# Development of a digital analysis system for a novel 3D culture-based colony formation to detect malignantly transformed cells in human cell-based therapeutic products

**DOI:** 10.1101/2025.05.08.651353

**Authors:** Shinji Kusakawa, Tingshu Yang, Rumi Sawada, Yoji Sato, Satoshi Yasuda

**Author notes:** Correspondence: Shinji Kusakawa, Division of Cell-Based Therapeutic Products, National Institute of Health Sciences, 3-25-26 Tonomachi, Kawasaki Ward, Kawasaki City, Kanagawa 210-9501, Japan., E-mail addresses. Correspondence: Satoshi Yasuda, Division of Cell-Based Therapeutic Products, National Institute of Health Sciences, 3-25-26 Tonomachi, Kawasaki Ward, Kawasaki City, Kanagawa 210-9501, Japan., E-mail addresses.

## Abstract

The presence of malignantly transformed cells in human cell-based therapeutic products (hCTPs) is a significant safety concern. Although such cellular impurities in hCTPs can be assessed by detecting anchorage-independent growth using conventional soft agar colony formation (SACF) assays, the sensitivity of these assays is often insufficient. To overcome this limitation, we previously developed a novel tumorigenicity-associated testing method, the digital SACF (D-SACF) assay, which combines a partitioned culture of test cells to concentrate target cells with colony detection via image analysis. However, conventional soft agar culture involves complicated operations, such as preparing multilayered culture media and temperature control, and further technical optimization is required for the widespread adoption of the D-SACF assay. In this study, we focused on a new culture system incorporating a three-dimensional (3D) culture method using a liquid medium containing the low-molecular-weight agar polymer LA717 in low-adhesion culture vessels. We initially confirmed conditions for the efficient high-density 3D culture of normal cells using LA717-supplemented medium in low-adhesion 96-well plates. Using human mesenchymal stem/stromal cells (MSCs) as a normal cell model and HeLa cells as a transformed cell model, we demonstrated that the new 3D culture system effectively maintained the dispersion of MSCs and prevented their aggregation, while transformed HeLa cells exhibited robust anchorage independence, thereby establishing the new liquid/low-molecular-weight agar colony formation (LACF) method as an alternative to SACF. Finally, by systematizing the digital analysis system for the LACF assay (D-LACF assay), which streamlines the overall workflow from the performance evaluation of the test method to product testing and result interpretation, the limitations of the conventional soft agar-based D-SACF assay were addressed and its practicality and utility were enhanced. This *in vitro* evaluation system is expected to provide a promising approach for improving the quality and safety of hCTPs.

## Introduction

In a three-dimensional (3D) culture environment without an adhesion substrate, normal cells typically undergo anoikis, a type of programmed cell death triggered by the loss of cell–matrix interactions [1,2]. In contrast, malignantly transformed cells exhibit anchorage-independent growth, allowing for their survival, proliferation, and colony formation under such conditions. This distinct phenotype is a hallmark of malignant transformation, and serves as a critical parameter for assessing cellular tumorigenicity [3]. The soft agar colony formation (SACF) assay is regarded as the gold standard for the *in vitro* assessment of cellular anchorage-independent growth [4–6]. Over the years, this assay has been refined to fulfill specific experimental needs, resulting in various modifications that enhance its practicality and sensitivity [7]. For example, the conventional SACF assay is insufficiently sensitive to evaluate the presence of transformed cells as cellular impurities in human cell-based therapeutic products (hCTPs), which consist of large numbers of cells [8]. To overcome this problem, we previously established the digital SACF (D-SACF) assay as an innovative test method that integrates soft agar culture, partitioned culture in multi-well plates, and image analysis to enable the digital detection of colonies derived from trace amounts of transformed cells within a population of normal cells [9]. Using an imaging cytometer for high-throughput image acquisition and automated analysis, this approach allows the efficient screening of large numbers of cells with high sensitivity to detect mixed transformed cells. In a previous study, the detection limit of the D-SACF assay, when evaluating its performance using human mesenchymal stem/stromal cells (MSCs) as the product model and HeLa cells as the transformed cell model, was recorded as one HeLa cell in 1×10^7^ MSCs, i.e., 0.00001% of the total number of cells. Furthermore, recent multisite studies have confirmed the high utility and feasibility of this test system [10]. Despite these advantages, the D-SACF assay presents practical challenges. As the number of wells and plates increases, preparing multilayered agar media with different concentrations and maintaining the required temperature becomes challenging. This procedural difficulty imposes a significant barrier to the feasibility and throughput for the large-scale application of the assay, underscoring the need for a more streamlined and optimized approach for the D-SACF assay.

To address these issues, we focused on a novel 3D culture system that utilizes a commercially available low-molecular-weight agar polymer LA717 (developed by Nissan Chemical Corporation, Tokyo, Japan)-supplemented medium combined with low-attachment culture vessels. Previous studies have demonstrated that using liquid culture media containing LA717 and cultivating cells in low-adhesion containers enables the even dispersion of cells throughout the culture medium, preventing adhesion and aggregation of neighboring cells while maintaining contact between daughter cells after division [11,12]. Thus, this system provides an easier means of establishing a 3D environment than conventional soft agar preparation, allows for a high-density cell culture, and may serve as a platform for evaluating anchorage-independent cell growth.

In this study, we used human MSCs as a model of normal cells and HeLa cells as a model of transformed cells to determine whether the novel liquid/low-molecular-weight agar colony formation (LACF) approach utilizing LA717 could effectively detect colony formation of malignantly transformed cells through anchorage-independent growth within the normal cell population. To overcome the limitations of the soft agar-based D-SACF assay and further improve its practicality and throughput, we optimized and systematized the protocol, ultimately establishing a digital analysis system for the LACF assay (D-LACF assay). Herein, we demonstrate how the D-LACF assay allows for the detection of malignantly transformed cells as tumorigenic impurities in hCTPs with high sensitivity and reproducibility, providing a promising new approach for the quality assessment of hCTPs.

## Materials and Methods

### Cells

All cell cultures were maintained in a humidified atmosphere with 5% CO_2_ and 95% air at 37°C. iCell Human iPSC-derived Mesenchymal Stem Cells (Lot 105305, Fujifilm Cellular Dynamics, Madison, WI, USA) were cultured using the human Mesenchymal Stem Cell Growth BulletKit Medium (Lonza, Basel, Switzerland). A human cervical cancer cell line, wild-type HeLa (CCL-2, Lot 61647128), was obtained from the American Type Culture Collection (Manassas, VA, USA). We obtained plasmids pC13N-iCAG.copGFP, pZT-C13-R1, and pZT-C13-L1 described by Cerbini *et al*. [13] from Addgene (Watertown, MA, USA) and introduced them into HeLa cells by electroporation. This approach enabled transcription activator-like effector nuclease (TALEN)-mediated green fluorescent protein (GFP) knock-in at the second intron of *CLYBLE* on chromosome 13, which is recognized as a safe harbor site. A stable GFP-expressing HeLa (HeLa-GFP) cell line was established for use in this study. Wild-type HeLa (HeLa-WT) and HeLa-GFP cells were cultured in Minimum Essential Medium (MEM; Sigma-Aldrich, St. Louis, MO, USA) supplemented with 10% fetal bovine serum (FBS, Sigma-Aldrich), 1% GlutaMAX (Thermo Fisher Scientific, Waltham, MA, USA), and 1% penicillin-streptomycin (Nacalai Tesque, Kyoto, Japan). MSCs and HeLa cells pre-labeled with CellVue Claret (CVC) dye (Sigma-Aldrich) were used in specific experiments following the manufacturer’s instructions.

### Novel LACF assay

We adopted a 3D culture method using a medium containing 0.03% (w/v) aqueous solution of LA717, which was supplied as a 1% stock solution under the trade name SphereMax (Nissan Chemical Corporation, Tokyo, Japan), based on previous research [11] by Abe-Fukasawa *et al*. This culture method facilitates straightforward 3D cell culture with a standard cell culture medium augmented by a fixed amount of LA717 (manufacturer’s recommended concentration: 0.03%) in low-adhesion culture vessels. In this study, cells were prepared in a prewarmed medium made up of a basal growth medium containing GlutaMAX-supplemented Dulbecco’s Modified Eagle Medium (Thermo Fisher Scientific), prewarmed dissolved LA717, 10% FBS, 1% MEM Non-Essential Amino Acid solution (Thermo Fisher Scientific), and 1% penicillin-streptomycin to achieve a final concentration of 0.03–0.06% LA717 (w/v). The cell suspensions were adjusted to a density of 15,000–600,000 cells/mL and dispensed into 96-well flat-bottom low-adhesion plates (PhenoPlate 384-well ULA-coated microplates, Revvity, Tokyo, Japan) at 100–400 μL (containing 10,000–60,000 cells) per well. These plates were incubated in a humidified atmosphere with 5% CO_2_ at 37°C for approximately 3 weeks (19–23 days) without any media changes or addition.

### Preparation of co-cultured cells: normal and reference transformed cells

In this study, MSCs were used as the normal cell model, whereas HeLa (-WT or -GFP) cells served as the reference transformed cell model. Samples were prepared by mixing these cells at a defined ratio, and the colony formation rate of individual transformed cells was evaluated under 3D culture conditions. To each well of a 96-well plate, 200 µL of medium supplemented with 0.03% LA717 was added, along with 20,000 MSCs and 0.125 to 1 HeLa cell. We employed a method in which the reference transformed cell suspension was initially prepared at a minimum concentration of 100 cells/mL using a procedure described later to minimize errors associated with repeated dilution steps. The required volume was then directly taken from this suspension and added to the MSC suspension, which had been preloaded into a reservoir. After thoroughly mixing, the suspensions were seeded into multiple wells. Additionally, when sampling a fixed volume from a low-concentration cell suspension, we considered the Poisson distribution of the actual cell count along with its 95% confidence interval (CI). The CI for the Poisson distribution was calculated using the poisson.test function in R (version 3.3.3, R Foundation for Statistical Computing, Vienna, Austria). For instance, when spiking a suspension expected to contain 100 cells, executing “poisson.test (100, conf.level = 0.95)” in R yields a 95% CI ranging from 81–122 cells (indicating a rounded-down or rounded-up value).

### Cell counting

The concentration of viable cells in the cell suspension was measured using NucleoCounter NC-202 Consistent Cell Counter (ChemoMetec, Cambridge, MA, USA) or Countess 3 FL Automated Cell Counter (Thermo Fisher Scientific). Based on these results, serial dilutions were performed to prepare suspensions of appropriate concentrations for each test. When preparing low-concentration HeLa cell suspensions at 100 cells/mL, a portion of the diluted suspension was used to reconfirm the concentration using the following method: 1.2 mL of the suspension was collected and incubated at 37°C for 15 min with 1 µg/mL Hoechst 33342 (Dojindo Laboratories, Kumamoto, Japan), 1 µg/mL propidium iodide (Dojindo Laboratories), and 1 µg/mL Calcein-AM (Dojindo Laboratories) (not added when counting HeLa-GFP), and then dispensed into 96 wells of a 384-well plate (Corning, Corning, NY, USA) at 10 µL each.

After centrifugation of the 384-well plate at 300 × *g* for 1 min, the presence or absence of viable cells in each well was analyzed by fluorescence imaging using the confocal imaging cytometer CellVoyager CQ1 (Yokogawa Electric, Musashino, Japan). Based on the number of negative wells (k) without cells out of the 96 wells, the average cell count (λ) per 10 µL was calculated using the formula λ = -*ln*(*k*), similar to determining the average copy number per partition in droplet digital PCR (ddPCR) [14]. The final concentration of the suspension was thus determined.

### Cell staining

After 3 weeks of incubation, the cells were stained with various reagents under the following conditions. The LACF assay medium (containing 0.03% LA717) with each staining dye was carefully added to each well of the plate at 20 µL. The plate was then incubated at 37°C and 5% CO2 for at least 1 h. The final concentrations of each staining reagent after addition were as follows: Hoechst 33342 at 1 µg/mL, Calcein-AM at 1 µM, LysoTracker Red (Thermo Fisher Scientific) at 50 nM, or MitoTracker Deep Red (Thermo Fisher Scientific) at 50 nM. The images of each well were captured using an imaging cytometer. In this study, we used two different imaging devices: the CellVoyager CQ1 (Yokogawa Electric) and the Cell3iMager duos2 (SCREEN Holdings, Kyoto, Japan). The former was primarily used for acquiring fluorescent images for analysis, whereas the latter was mainly employed to obtain clear brightfield images of the entire well. When fixing samples stained with reagents other than Calcein-AM, 15 µL of 16% paraformaldehyde (PFA) solution (Muto Pure Chemicals, Tokyo, Japan) prewarmed to 37°C was added to each well (with a final PFA concentration of approximately 1%). The samples were kept in the dark and allowed to stand overnight at 37°C. Subsequently, the samples were stored in the dark at room temperature. Fixed samples were imaged within 2 weeks of this procedure.

### Image analysis

An imaging cytometer was used to capture images of each well. Excitation light and fluorescence filters corresponding to the fluorescent proteins expressed in the cells or the added fluorescent reagents were selected and multichannel images of the entire well, including bright-field images, were obtained. Furthermore, multiple Z-stack images were acquired, and images focused on the cell colonies were selected. Automated image analysis was conducted for each well using an application integrated into the analyzer, employing an analysis script to segment colony features based on size, circularity, and fluorescence intensity criteria. In this study, through image analysis, the cell survival status was determined based on the expression of fluorescent proteins or the staining reaction of cells to fluorescent reagents. HeLa cell-derived colonies were then defined and detected based on their size, shape, and circularity. However, in some cases, debris and other objects were detected as false-positive objects in the image analysis; therefore, the presence or absence of colonies in each well was ultimately confirmed by visual inspection based on bright-field images.

### Calculation of the colony-forming efficiency (CFE) of reference transformed cells

Calculations were performed to determine the CFE (x) of reference transformed cells based on the following assumptions:

1. Cells are assumed to be individually dispersed in the suspension without aggregation.
2. Transformed cells seeded into a fixed number of wells (n) are distributed across the wells following a Poisson distribution.
3. Under 3D culture conditions, a single transformed cell in a well of a 96-well plate proliferates and forms a colony with a specific probability (0≤x≤1).
4. When multiple transformed cells are seeded into the same well, they each form colonies independently with the same probability (x), without influencing one another.

Using these assumptions, the relationship among the number of fractionated wells (n), the average number of transformed cells per well (λ), the total number of wells with colonies (y), and the probability of colony formation (x) is described by the following equation (see Supplementary Figures 1 and 2 for detailed mathematical derivation and application examples):

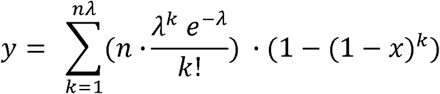

To determine x, the upper limit of k was estimated based on the actual number of fractionated wells (n) and the average number of transformed cells per well (λ). As shown in Supplementary Figure 1, the value of k was set to a level that ensures the number of wells with colonies is not zero, regardless of the CFE, based on the seeding conditions (particularly n and λ) of the reference transformed cells. Calculations for x were performed using the Goal Seek function (What-If Analysis) in Microsoft Excel (version 16.96; Microsoft, Redmond, WA, USA) to obtain approximate results.

### Statistical analysis

Statistical analyses were performed using Prism 10 software (version 10.3.0; GraphPad Software, San Diego, CA, USA). Statistical significance was defined as *P* < 0.05. The effects of culture conditions and independent test trials on the mean CFE of HeLa-GFP cells under different culture conditions were evaluated by one-way repeated measures analysis of variance (ANOVA). CFE values are presented as the mean ± standard deviation or with their corresponding 95% CI.

## Results

### LA717 enables stable and dispersed 3D MSC cultures at a high cell density

To evaluate the effect of LA717 on the high-density 3D culture of MSCs, cells were seeded at various densities (10,000–60,000 cells/well) in low-adhesion culture plates. In the absence of LA717, MSCs exhibited rapid aggregation, forming dense cell clusters within the first day of culture, regardless of the initial seeding density (Figure 1, left panel). This aggregation persisted and became more pronounced over time, especially in the wells with higher cell densities. In contrast, the addition of 0.03% LA717 to the culture medium significantly suppressed MSC aggregation. The cells remained largely dispersed across all seeding densities even after 14 days of culture (Figure 1, right panel). Although slight aggregation was observed at higher cell densities (50,000 and 60,000 cells/well), these aggregates were transient and shrank over time, indicating that LA717 effectively promoted a stable dispersed state of MSCs under high-density conditions.

**Figure 1.**
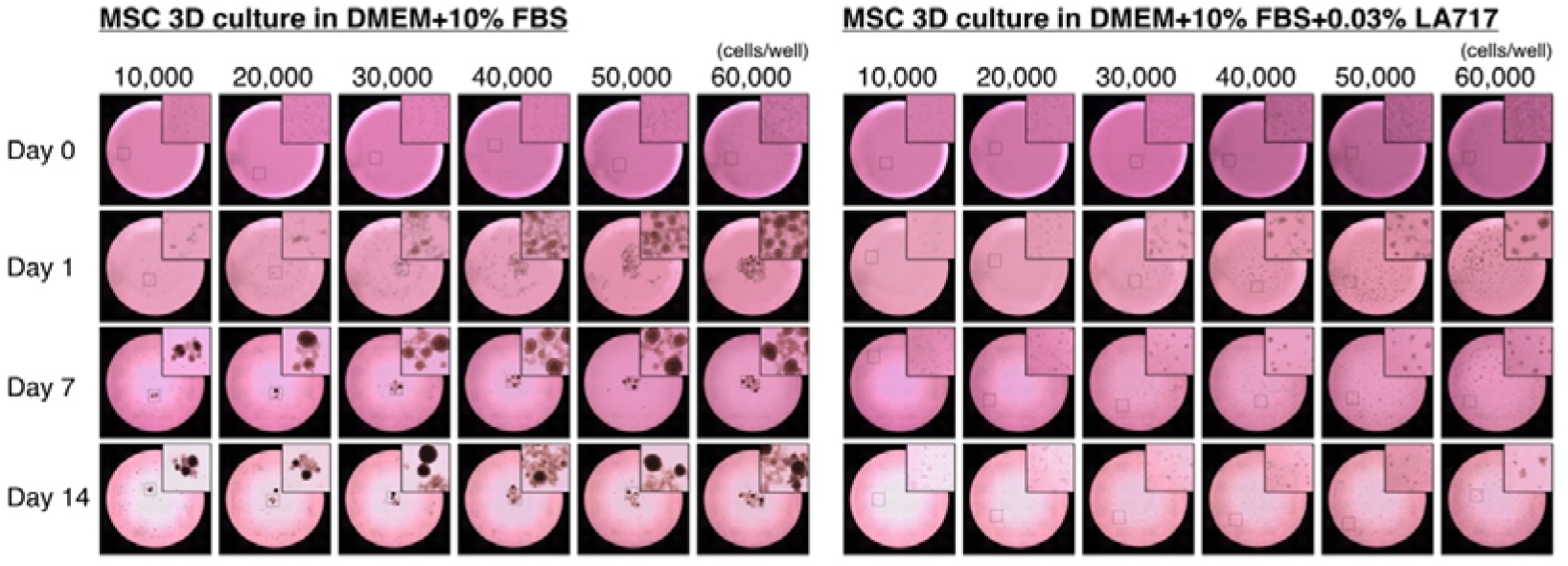
LA717 prevents cell aggregation in high-density three-dimensional (3D) cultures of human mesenchymal stem/stromal cells (MSCs). Representative whole-well images showing the time course of MSC cultures over 14 days in low-adhesion 96-well plates under two different 3D culture conditions. Insets show magnified views of the boxed areas to highlight cell aggregation or distribution. MSCs were seeded at densities ranging from 10,000 to 60,000 cells/well in Dulbecco’s Modified Eagle Medium (DMEM) supplemented with 10% fetal bovine serum (FBS; left) or DMEM supplemented with 10% FBS and 0.03% LA717 (right; developed by Nissan Chemical Corporation, Tokyo, Japan). In the absence of LA717, MSCs aggregated rapidly, particularly at higher seeding densities, and formed dense clusters by day 1 that persisted throughout the culture period. In contrast, the addition of 0.03% LA717 effectively suppressed cell aggregation, maintaining a dispersed distribution of MSCs across all densities over the entire 14-day period. All images were acquired using the Cell3iMager duos2 (SCREEN Holdings, Kyoto, Japan). The inner diameter of each well was 6.4 mm.

### Culture conditions to optimize high-density 3D MSC culture

In this study, we investigated the effects of LA717 concentration and medium volume on the aggregation of MSCs in cultures in 3D using high-density cultures of 60,000 cells/well. MSCs cultured without LA717 showed significant aggregation regardless of the medium volume. However, the addition of LA717 effectively suppressed aggregation in a concentration-dependent manner, with higher concentrations of LA717 (0.04–0.06%) showing more pronounced effects (Figure 2). Furthermore, large medium volumes (300–400 µL) were associated with a greater decrease in MSC aggregation, even at low LA717 concentrations (0.03–0.04%). These results suggest that both the LA717 concentration and medium volume play crucial roles in preventing MSC aggregation in high-density 3D culture.

**Figure 2.**
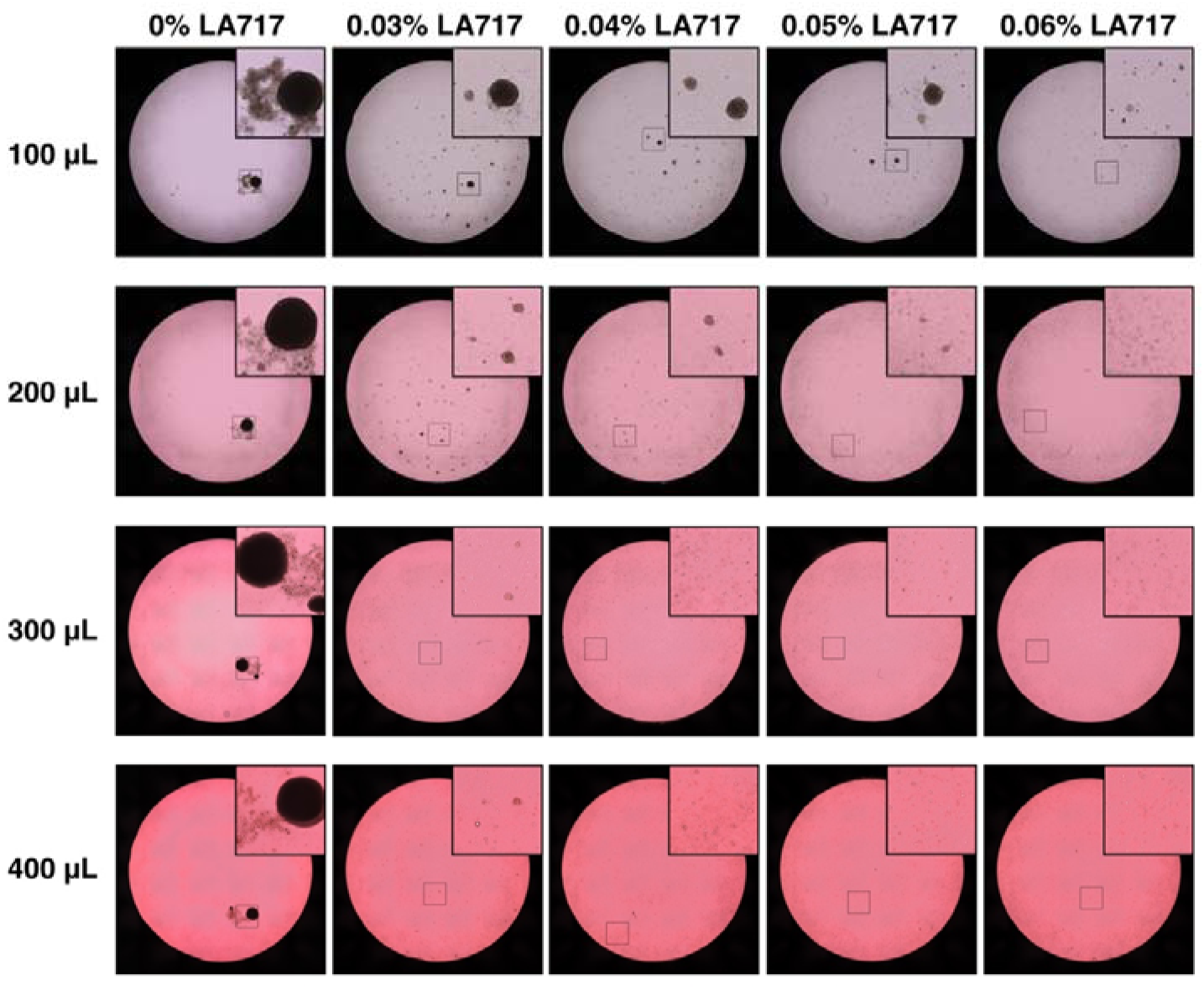
LA717 suppresses MSC aggregation in a concentration- and volume-dependent manner in 3D culture. Representative whole-well images showing the effects of LA717 concentration and culture medium volume on MSC aggregation after 14 days of 3D culture. Insets show magnified views of the boxed areas to highlight cell aggregation or distribution. MSCs (60,000 cells/well) were cultured in low-adhesion 96-well plates with DMEM + 10% FBS containing varying concentrations of LA717 (0–0.06%) and medium volumes (100, 200, 300, and 400 µL). In the absence of LA717, MSCs aggregated extensively regardless of the medium volume. The addition of LA717 progressively suppressed aggregation in a concentration-dependent manner, with higher concentrations (≥0.04%) and greater medium volumes (≥300 µL) showing improved dispersion. These findings suggested that both LA717 concentration and medium volume are important parameters for achieving a stable, high-density 3D culture of MSCs. All images were acquired using the Cell3iMager duos2 (SCREEN Holdings). The inner diameter of each well was 6.4 mm.

### A single HeLa cell co-cultured with MSCs in LA717-supplemented medium proliferated in an anchorage-independent manner and formed a colony

Next, we examined the behavior of MSCs and a single HeLa-GFP cell co-cultured in a 3D environment with medium supplemented with 0.03% LA717 (200 µL per well). All cells were pre-labeled with the CVC dye for fluorescence imaging (Figure 3). Bright-field and CVC fluorescence (pseudo-colored in magenta) images demonstrated that MSCs remained dispersed and showed no anchorage-independent growth under this 3D culture condition. Minor initial aggregation of MSCs was observed, which resolved over time. In contrast, the single HeLa-GFP cell exhibited anchorage-independent growth, doubled within 24 h, and continued to proliferate, ultimately forming a distinct colony. GFP fluorescence highlighted the dynamic HeLa cell colony formation process over 15 days. However, the fluorescence of the CVC dye in colonies derived from HeLa-GFP cells almost disappeared, likely due to the continuous dilution of the fluorescent label as the cells divided. These results suggest that the 3D culture system utilizing LA717 effectively distinguishes the proliferative capacity of abnormally transformed cells, such as HeLa cells, from that of normal MSCs. The images in Figure 3 show a section at each time point when time-lapse imaging (360 h, imaging at 1 h intervals) was performed, and the video showing all the images for each time point stitched together is shown in the Supplementary Movie.

**Figure 3.**
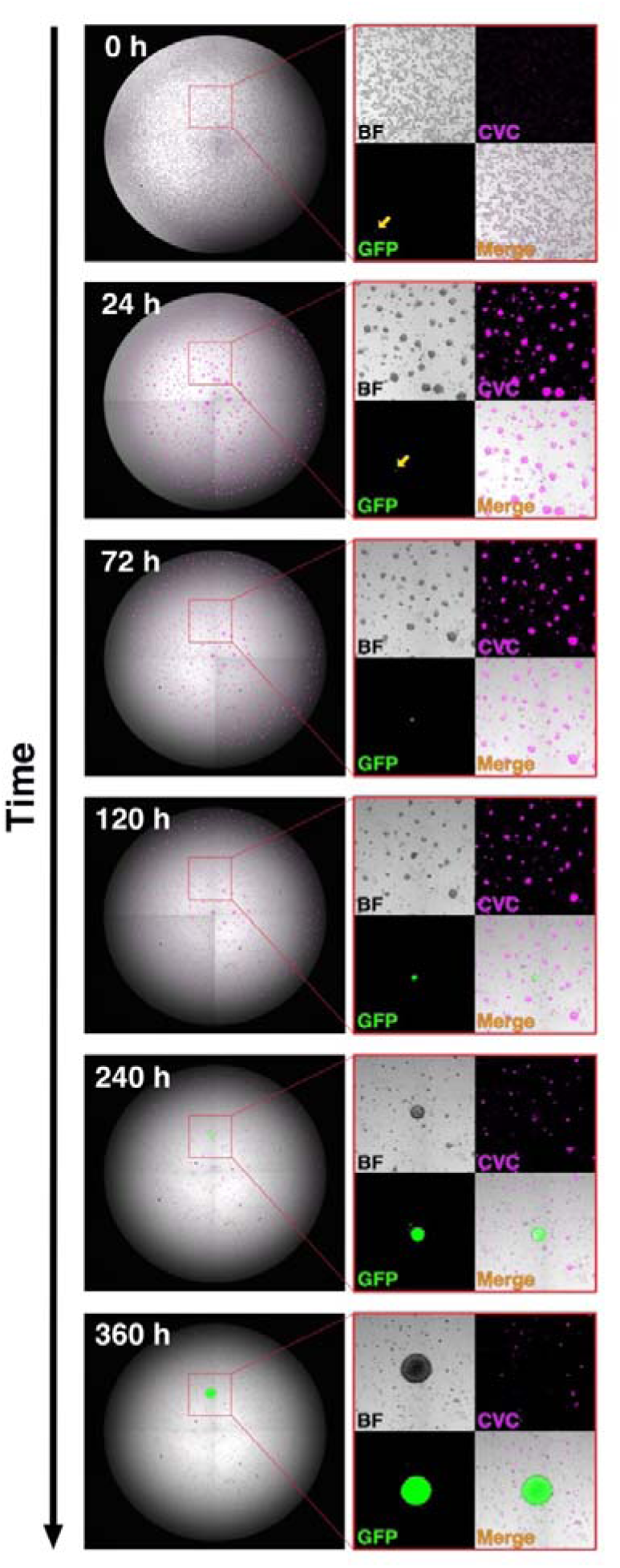
Transformed HeLa-green fluorescent protein (GFP) cells exhibit anchorage-independent proliferation and colony formation in a 3D culture environment with 0.03% LA717, unlike surrounding MSCs. Representative images extracted from time-lapse imaging performed every hour for 360 h (15 days), showing the behavior of a single HeLa-GFP cell co-cultured with 20,000 MSCs in 200 µL of LA717 (0.03%)-supplemented medium in a low-adhesion 96-well plate. All cells were pre-labeled with CellVue Claret (CVC; pseudo-colored magenta) for fluorescence visualization. Bright-field (BF), GFP, CVC, and merged images at six time points revealed that, while MSCs remained dispersed and did not proliferate, the single HeLa-GFP cell consistently proliferated in an anchorage-independent manner, ultimately forming a distinct colony. The GFP signal expanded over time, highlighting the dynamic growth of the transformed cells. A stitched movie of all time points is provided in the Supplementary Movie. Images were acquired using a CellVoyager CQ1 (Yokogawa Electric). The inner diameter of each well was 6.4 mm.

### Single HeLa cells consistently formed colonies with a steady efficiency across different 3D culture conditions in the presence of LA717

We previously established a method for calculating the CFE of reference transformed cells, which serves as an indicator of the detection performance of transformed cells in the D-SACF assay (an approach that allows the efficient analysis of a large number of cells). Conventionally, the CFE is determined based on the total number of counted colonies obtained from multi-well image analysis. However, in this study, we developed and implemented a new approach whereby the CFE is calculated based on the total number of wells containing at least one colony rather than the number of counted colonies. Using this method, we assessed the CFE of HeLa-GFP cells under various culture conditions.

First, we evaluated the CFE of HeLa-GFP cells co-cultured with MSCs in a medium containing LA717 at varying concentrations (0.03–0.06%). We seeded cells into a total of 240 wells (60 wells × 4 plates) under conditions where each well had 200 µL of medium with 20,000 MSCs and an average of 0.5 HeLa-GFP cells (i.e., λ = 0.5 cells/well). The cells were then co-cultured in a 3D environment, and colony analysis was performed after 3 weeks. The results of three independent runs for calculating the CFE of HeLa-GFP cells at four different concentrations of LA717 (0.03%, 0.04%, 0.05%, and 0.06%) are shown in Figure 4a. The data were analyzed using one-way repeated measures ANOVA, which revealed no significant effects of LA717 concentration on the CFE of HeLa-GFP cells. These findings indicate that the CFE of HeLa-GFP cells remained stable regardless of the LA717 concentration. This stability suggests that LA717 did not influence the anchorage-independent growth capacity of HeLa-GFP cells within the tested concentration range.

**Figure 4.**
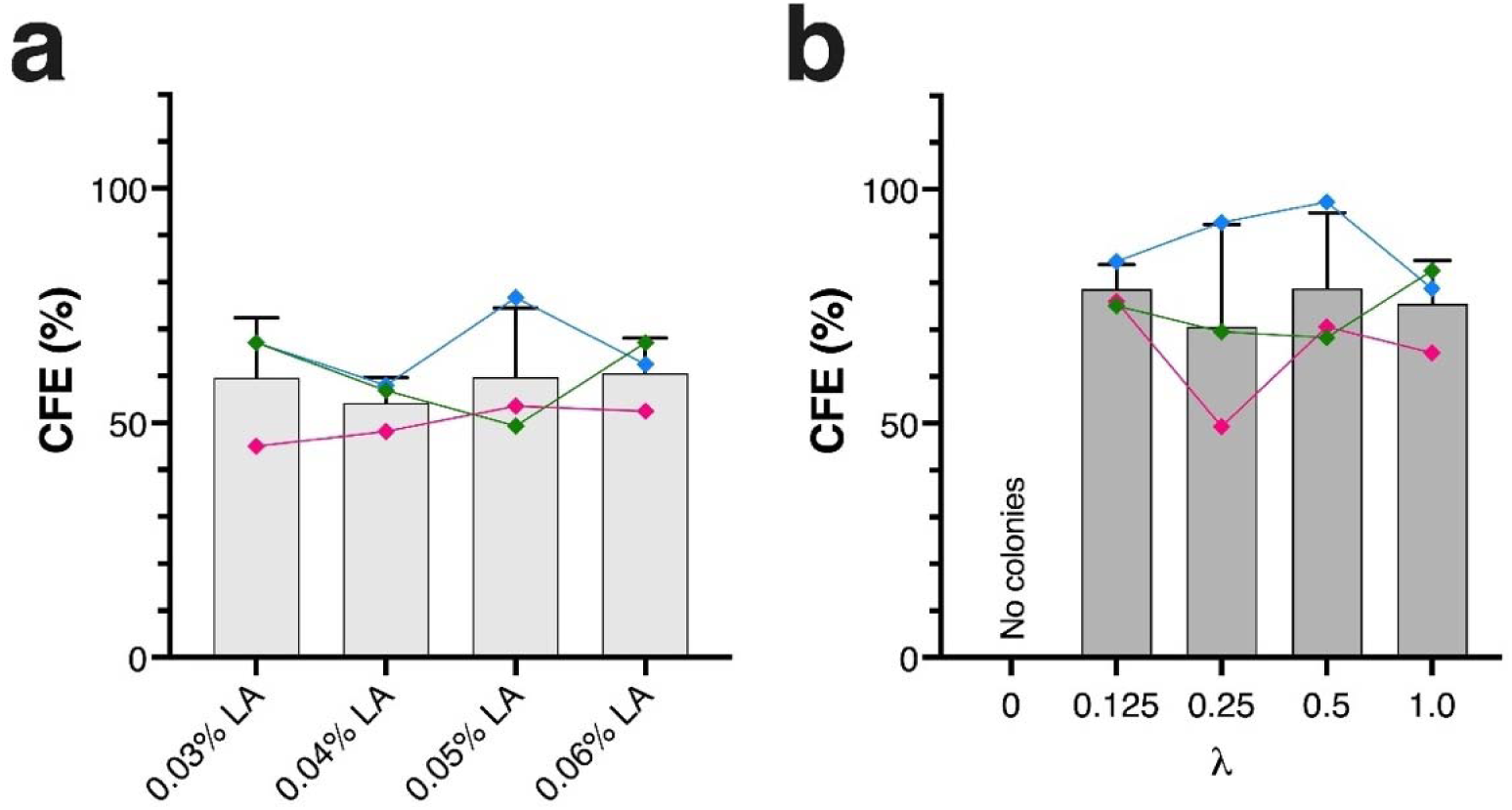
The colony-forming efficiency (CFE) of HeLa-GFP cells co-cultured with MSCs in LA717-supplemented medium under various culture conditions. **(a)** The CFE of HeLa-GFP cells was evaluated under four different concentrations of LA717 (0.03%, 0.04%, 0.05%, and 0.06%) in 200 µL of culture medium per well. HeLa-GFP cells (0.5 cells/well on average) were co-cultured with 20,000 MSCs per well in a 96-well plate (*n* = 240 wells per condition) for 3 weeks. CFE was calculated based on the number of wells containing colonies derived from HeLa-GFP cells, as determined by image analysis. Bar graphs represent the mean and standard deviation (SD) values from three independent experiments, and the colored lines show the individual results from each experiment. **(b)** The CFE of HeLa-GFP cells was evaluated under varying cell input amounts (λ = 0.125, 0.25, 0.5, and 1.0) in 200 µL of medium containing 0.03% LA717 and 20,000 MSCs per well. HeLa-GFP cells were seeded into 840, 420, 240, and 120 wells, respectively, and cultured for 3 weeks. The CFE was calculated from the number of wells containing colonies detected by image analysis. Bars represent the mean and SD values of three independent experiments, and colored lines indicate the individual results. No colony formation was observed under control conditions (λ = 0; i.e., MSCs only, 60 wells). Detailed numerical data for each condition in **(a)** and **(b)** are provided in Supplementary Tables 1 and 2, respectively. The equation used to calculate CFE in **(a)** and **(b)** is described in Supplementary Figures 1 and 2, respectively. LA, LA717.

Next, to evaluate the impact of seeding conditions on the CFE of HeLa-GFP cells, we conducted three independent runs under fixed culture conditions: 200[µL of the medium containing 0.03% LA717 and 20,000 MSCs per well. We varied the seeding ratio of HeLa-GFP cells (λ = 0–1.0 cells/well) and the total number of seeded wells (60–840 wells). After 3 weeks of culture, the CFE was analyzed under each condition. The results showed that the CFE of HeLa-GFP cells remained consistent across all tested seeding conditions (Figure 4b). Statistical analysis using one-way repeated measures ANOVA revealed no significant effect of the HeLa-GFP seeding ratio on the CFE. These findings indicate that under the examined conditions, the evaluation of CFE of reference transformed cells is robust and reliable. Importantly, under control conditions (λ = 0; MSCs only, 60 wells), no colony formation was observed, confirming the specificity of colony detection for HeLa-GFP cells.

Supplementary Tables 1 and 2 present the CFE of the reference transformed cells under each culture condition, calculated using two distinct methods: (1) based on the total number of wells containing at least one colony, as applied in this study, and (2) based on the conventional method of counting the total number of colonies. The results from both methods were consistent, indicating that the CFE of the reference transformed cells remained nearly identical regardless of the calculation method employed.

### Improved visualization and detection of colonies using fluorescence staining in LA717-based 3D culture

To enhance the efficiency and accuracy of colony detection through the image analysis of whole-well images in a 96-well plate, we explored the application of fluorescence staining in cultured samples. Based on our previous studies, we evaluated nuclear, mitochondrial, and lysosomal fluorescent labeling for this purpose. Figure 5a shows representative images of a cell colony observed after co-culturing 0.5 HeLa-GFP cells with 20,000 MSCs per well in a medium supplemented with 0.03% LA717 for 3 weeks. The presence of GFP fluorescence in the cell aggregates identified in the bright-field images confirmed that these colonies originated from HeLa-GFP cells. The addition of fluorescent dyes specific to the nucleus, mitochondria, and lysosomes during cell culture enabled the organelle staining of live cells, leading to clear fluorescent labeling of HeLa-GFP-derived colonies. Among the dyes used, LysoTracker and MitoTracker are selective for live cells, which indicated that the colonies consisted of viable cells. Figure 5b shows colonies formed by HeLa-WT cells under the same culture conditions. Live staining with Calcein-AM resulted in strong fluorescence signals, confirming that the colonies were composed of viable cells. Based on the findings from HeLa-GFP cells, the colonies observed under these conditions were likely derived from HeLa-WT cells.

**Figure 5.**
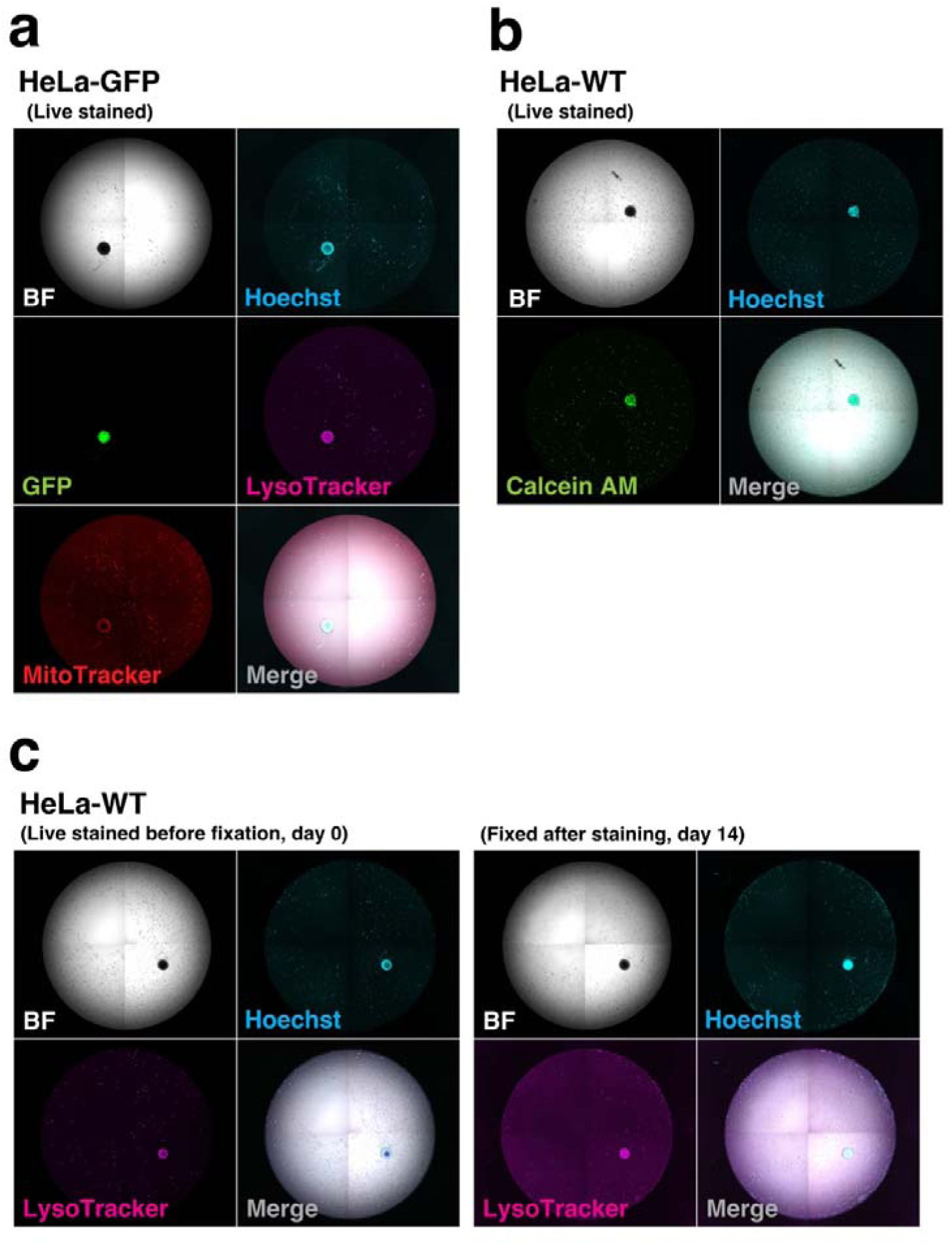
Fluorescence-based detection of colonies formed by single HeLa cells co-cultured with MSCs in LA717-supplemented medium. Representative images of colonies formed by a single HeLa-GFP or wild-type HeLa cell co-cultured with 20,000 MSCs per well for 3 weeks in 0.03% LA717-supplemented medium (200 µL/well). Fluorescent staining markedly enhanced the accuracy of colony detection in image analysis. Images were acquired using the CellVoyager CQ1 confocal imaging cytometer (Yokogawa Electric). **(a)** A colony derived from a single HeLa-GFP cell. Live staining was performed by adding Hoechst 33342 (nucleus), MitoTracker Deep Red (mitochondria), or LysoTracker Red (lysosomes) to the culture medium and incubating the cells for a defined period. The GFP signal confirmed that the colony originated from HeLa-GFP cells, which was also clearly visualized with each dye. **(b)** A colony derived from a single wild-type HeLa cell. Live staining with Hoechst 33342 and Calcein-AM revealed strong fluorescence signals, indicating colony viability. **(c)** (Left) A wild-type live HeLa colony stained with Hoechst 33342 and LysoTracker Red. (Right) The same colony was imaged 14 days after fixation with 1% paraformaldehyde (PFA). Fluorescence signals from Hoechst and LysoTracker staining were retained, indicating that this dye combination is compatible with fixation and long-term storage, allowing flexible scheduling of image acquisition and analysis. The inner diameter of each well was 6.4 mm.

Fluorescent staining has proven useful for image-based colony detection. To enhance its applicability, we further investigated fixation methods for stained cells. As shown in Figure 5c, careful temperature management during fixation allowed fluorescence signals from LysoTracker and Hoechst staining (left images show live colony staining before fixation) to be preserved for at least 14 days (right images show the fixed colony stained with LysoTracker and Hoechst dyes). This approach increases the flexibility of the analytical process by enabling imaging-based colony evaluation over an extended period, thereby improving the practicality of the method.

These fluorescent labeling techniques enabled clear visualization of colonies derived from transformed cells in the LA717-based 3D culture environment. Consequently, similar to the previous D-SACF assay, detecting transformed cell-derived colonies and evaluating the CFE of reference cells through image analysis became readily achievable.

## Discussion

A previous study conducted by the developers of LA717 [11] demonstrated that culturing cells in low-attachment vessels with LA717-supplemented medium ensures uniform cell dispersion, prevents adhesion and aggregation, and maintains contact between newly divided cells while minimizing medium convection. These properties allow cells and spheroids to stay in place and form the foundation of a new 3D culture system. Building on this, we showed that MSCs, which typically form aggregates at high densities, remained stably dispersed without proliferating in the LA717-containing medium, supporting its use for scalable and uniform MSC culture. In contrast, HeLa cells exhibited anchorage-independent growth from single cells and consistently formed colonies under the same conditions. Fluorescent labeling enhanced the visualization of colonies from transformed cells, making image-based colony detection easier and supporting CFE evaluation with reference cells such as HeLa cells. The LACF assay, developed using this 3D culture system with LA717, demonstrated practical advantages over the SACF assay in terms of operability and analysis (Supplementary Table 3). We further established the D-LACF assay to digitally measure the proportion of transformed cells in hCTPs by detecting single-cell-derived colonies (Figure 6).

**Figure 6.**
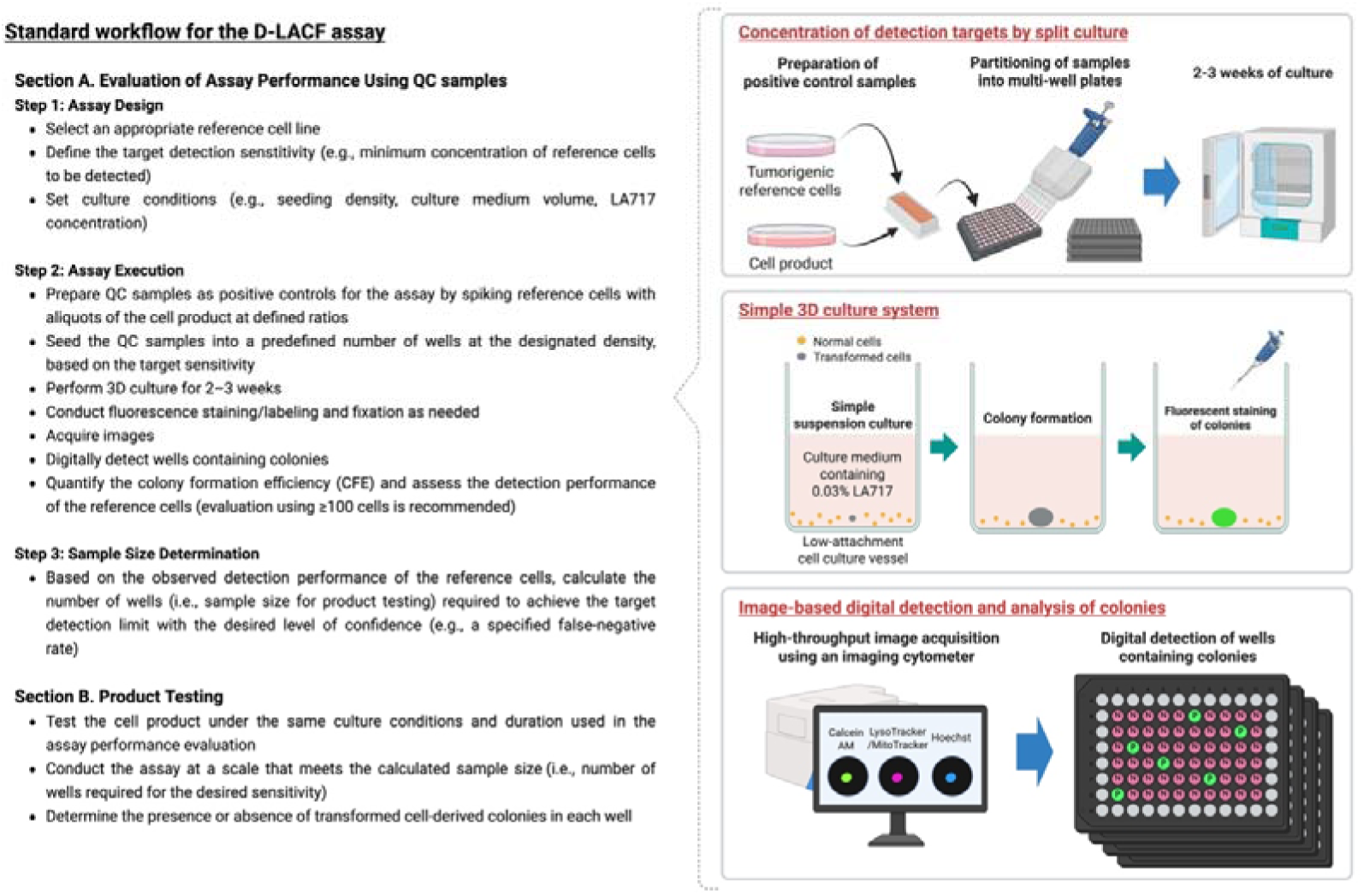
Schematic overview of the digital liquid/low-molecular-weight agar colony formation (D-LACF) assay. The D-LACF assay can be used to detect transformed cells by combining LA717-based 3D culture with image□based digital analysis. Section A illustrates the evaluation of the assay performance using quality control (QC) samples: (1) assay design, (2) assay execution including split culture, 3D growth in LA717-supplemented medium, fluorescent labeling, and high-throughput colony detection, and (3) determination of the sample size based on the observed detection performance. Section B presents product testing conducted under the same culture conditions but with the predetermined sample size (number of wells) calculated in Section A; each well can then be classified as positive (P) or negative (N) for transformed cell-derived colonies. This figure was created with BioRender.com.

Quality control (QC) of the assay itself, using MSCs as a cell product and HeLa cells as the reference, requires a 3-week co-culture period, followed by staining, fixation, imaging, and analysis, which is typically completed within the same day or a few days. Based on this workflow, we apply product testing under the same conditions. In our prior D-SACF study, we developed a method to calculate the required sample size (i.e., the number of partitioned wells) to detect at least one colony at a predefined false-negative rate (FNR), based on the CFE of the reference cells [10]. This approach allows for the theoretical estimation of the assay scale needed to achieve a target detection sensitivity and FNR for QC samples containing transformed cells at a known frequency. It also enables the prediction of the probability of detecting at least one colony when using QC samples, given the pre-calculated sample size. In product testing, applying the same seeding conditions and number of wells helps determine whether transformed cells comparable to the reference are present and if their proportion exceeds the predefined detection limit—information that can be considered a quality attribute of the product. This method is equally applicable to the D-LACF assay, supporting rational, quantitative test design for assessing transformed cells in hCTPs under defined conditions. By routinely monitoring assay performance, especially through system suitability testing with reference cells, the D-LACF assay can be used for lot-release testing of cell products. Its improved feasibility and scalability over the D-SACF assay support large-scale screening of malignantly transformed cells in hCTPs.

The growth in low attachment assay (GILA) is another *in vitro* 3D culture system for transformation assessment using only standard medium and low-adhesion vessels [15,16]. It reportedly detects gene mutation-induced transformation with sensitivity comparable to SACF assay [17]. The possibility of multiple assay options is advantageous as it enables the selection of the most appropriate method based on various requirements. The standard GILA protocol recommends seeding several hundred to several thousand cells per well in a 96-well plate. However, in this study, when MSCs were seeded at 10,000 cells/well in a conventional culture medium, pronounced cell aggregation was observed. In contrast, the LA717-supplemented medium effectively suppressed MSC aggregation under the same seeding conditions. These results suggest that for quality assessment of hCTPs composed of a large number of cells—particularly for detecting a trace amount of transformed cells—GILA may not suitable for this purpose due to its limitations under high-density culture conditions. Instead, the LACF assay, which enables high-density culture by modifying the culture medium composition, is currently the most practical and effective method for such applications.

The D-SACF/D-LACF assay share core principles with ddPCR and is structured around three main components: (1) Fractionation of test cells into multi-well plates for 3D culture to enrich target transformed cells and improve the signal-to-noise ratio for image analysis; (2) Use of high-throughput imaging systems for efficient multi-well analysis; and (3) Digital detection of colony presence or absence in each well to estimate the proportion of transformed cells. Similar to ddPCR, increasing the number of partitions theoretically improves the detection of rare transformed cells, enhancing sensitivity and lowering the limit of detection (LOD). However, in the D-LACF assay, each well must receive a fixed number of total cells (including both normal and transformed cells), followed by 3D culture and colony assessment. This imposes constraints on optimal culture volume and cell density per well. Although switching from 96- to 384-well plates could improve resolution and throughput, challenges such as reduced seeding volume, higher risk of medium evaporation, and variations in cell growth might outweigh these benefits. Nonetheless, optimizing culture formats, including plate types, remains a crucial area of research for future assay automation. A key difference from ddPCR is that while a target sequence in ddPCR is almost always amplified, making the partition positive, not all transformed cells in D-SACF/D-LACF necessarily form colonies. Thus, estimating their presence ratio based solely on colony detection is difficult. This reflects a key challenge in detecting transformed cells with unknown properties in a product. To address this, the D-SACF/D-LACF assay uses a known transformed cell line, such as HeLa, as a reference and positive control. Its CFE is pre-estimated and used as a performance indicator for the assay.

In this study, we improved part of the performance evaluation process from the d-SACF assay. While assessing the CFE of reference cells, a key performance indicator, we focused on two operational challenges: the difficulty of preparing low-concentration cell suspensions and the CFE variability potentially caused by the cell spiking process. To improve the accuracy and consistency of the CFE assessment, we optimized the protocol for these steps. We refined the counting process by staining cells in 384-well plates and using image-based counting, which partially automates the workflow. We also incorporated Poisson distribution modeling and its 95% CI to account for sampling error during spiking, enabling a more precise definition of the expected range in performance metrics and enhancing analytical accuracy. Additionally, we developed a simplified way for calculating CFE. Since tracking cell-to-colony conversion precisely is impractical, we initially calculated the CFE as the ratio of the estimated number of seeded cells to the total number of colonies. However, image analysis is often complicated by overlapping or touching colonies, making exact counting difficult. To address this, we developed an alternative approach: estimating CFE based on the number of wells containing at least one colony. This method requires only binary detection of colonies (presence or absence), significantly simplifying image analysis. CFE values obtained with both methods were consistent and stable across various conditions, including different LA717 concentrations and HeLa-GFP seeding densities. These results demonstrate the robustness and reliability of the proposed approach, which we have adopted as the standard.

Colony imaging and analysis in the D-LACF assays can be performed using a standard fluorescence imaging system with multi-well scanning capabilities. In addition to previously used systems (IN Cell Analyzer 2000 [Cytiva], Cellomics ArrayScan [Thermo Fisher Scientific], and CellVoyager CQ1 [Yokogawa Electric]), this study utilized the Cell3iMager duos2 (SCREEN Holdings), which demonstrated comparable feasibility. To facilitate the widespread implementation of the D-LACF assays across laboratories, the minimum requirements for an appropriate imaging system are outlined in Supplementary Table 4. Image analysis remains a limiting factor. False positives can arise from fluorescent debris or cell aggregates, which are common with dyes like Hoechst 33342 or LysoTracker, or from insufficient mixing of the suspension. We addressed this by pre-labeling cell membranes with CVC, which allowed for the distinction between aggregates and proliferative colonies. Aggregates showed stable fluorescence but no growth; true colonies exhibited decreasing signal due to division. Incorporating metrics like total or mean fluorescence intensity could improve identification accuracy. False negatives are actual colonies that go unnoticed. Our current protocol focuses on decreasing false negatives, even if it causes more false positives. Therefore, we use permissive image analysis settings, followed by manual review of fluorescence and bright-field images to exclude artifacts through a two-step verification process. To further enhance accuracy and scalability, advanced techniques using more specific fluorescent dyes or artificial intelligence (AI)-based label-free analysis will be important. Such methods could decrease both false positives and false negatives, making the D-LACF assay more robust and dependable.

In product cell testing, the ideal outcome is that no colonies are detected. However, if colony-like structures are observed and their status as true or false positives is uncertain, the LACF assay offers a complementary option that the conventional SACF assay lacks. Because the LA717-based 3D culture system uses a liquid medium, suspect cell aggregates can be easily isolated and retrieved with a micropipette. The collected cells can then be sub-cultured to determine whether they proliferate, allowing for further characterization to establish whether they are truly malignant. Moreover, a detailed analysis of any unexpectedly tumorigenic cells may provide new opportunities to investigate the mechanisms of cellular transformation and develop strategies for its suppression, thereby widening the research applications of this novel 3D culture platform.

To facilitate broader adoption and practical use of the D-LACF assay, a systematic, stepwise approach—including inter-laboratory validation and standardization—is essential. Each participating facility should initially verify the assay’s reproducibility and feasibility using a standardized core protocol. We recommend referring to Supplementary Table 5, which complements Figure 6 and highlights key considerations and practical tips for implementing the D-LACF assay across laboratories. Overall, these efforts will help ensure the assay’s reliability and support its international standardization.

Method validation should then be conducted according to ICH Q2(R1) guidelines [19]. Since the D-LACF assay acts as a “limit test” within the “purity testing” category of QC, validation should mainly emphasize specificity and the LOD. In this study, we developed the D-LACF platform and demonstrated its feasibility using model cells: an iPSC-derived MSC line and HeLa cells. However, to assess specificity, it is necessary to evaluate additional tumor-derived cell lines and confirm that their colony formation can be detected. Although previous work showed that several cancer cell lines can grow in LA717-supplemented 3D culture [11], their CFEs have not been systematically characterized.

As a preliminary study, we evaluated the performance of the D-LACF assay using HEK-293 cells, a tumorigenic cell line different from HeLa (Supplementary Figure 3). Although the observed CFE of HEK-293 cells was lower than that of HeLa, consistent colony formation was still detected. These findings suggest the potential usefulness of the D-LACF assay for detecting various transformed cells. Further studies with different tumorigenic cell lines will be needed to assess their colony-forming capabilities, along with quantitative evaluations of spike-in concentrations and detection accuracy.

In the earlier D-SACF study, multiple MSC lots showed no lot-dependent effect on HeLa colony formation [10]. In this study, the iPSC-derived MSC line was chosen for its high proliferative capacity and consistent stock availability. Both D-SACF and D-LACF assays, which are based on similar principles, consistently showed that MSCs have no inhibitory effects on colony formation by HeLa cells. However, donor variability among MSCs has not been systematically investigated. Future studies should incorporate additional MSC lots, including clinical-grade cell batches, to assess assay robustness. Additionally, the applicability of the D-LACF assay to hCTP models beyond MSCs—such as neural or cardiac cell-based products—should be explored. For such validation, harmonizing reagents, cell sources, culture conditions, and assay design is critical. Inter-laboratory studies will be essential to confirm the reproducibility of key metrics, such as CFE.

In summary, the D-LACF workflow developed here can serve as a platform for designing QC protocols for various hCTPs. A key benefit of the assay is its flexibility—test parameters such as seeding density, well number, and LA717 concentration can be adjusted based on product characteristics. The D-LACF assay is anticipated to improve the quality and safety of hCTPs, supporting the development of more reliable therapies.

## Conclusion

In this study, we established the D-LACF assay as a practical and sensitive platform for detecting trace amounts of malignantly transformed cells in hCTPs. By combining a novel LA717-based 3D culture system with digital image analysis and refined evaluation protocols, the D-LACF assay offers improved feasibility, accuracy, and scalability compared with conventional methods. These findings provide a foundation for the broader implementation of the D-LACF assay in quality testing frameworks and contribute to enhancing the safety and reliability of future hCTPs. Moving forward, continued optimization and inter-laboratory validation of this assay will be crucial for its practical implementation and international standardization as a QC tool for advanced cell-based therapies.

## Author Contributions

Study conception and design: SK and SY. Data acquisition: SK and TY. Analysis and interpretation of data: All authors. Manuscript drafting and revision: SK and SY. All the authors have approved the final manuscript.

## Acknowledgments

We thank Keiichiro Otsuka and Taito Nishino from Nissan Chemical Corporation for their help and support in establishing the 3D culture method using LA717. We thank Atsune Kamada and Prof. Dr. Mitsutoshi Satoh of Meiji Pharmaceutical University for generating the preliminary data that laid the foundation for this study. We also express our gratitude to Takashi Yoshiura and Takayoshi Matsubara from Yokogawa Electric for their dedicated efforts in supporting and coordinating image acquisition and analysis with CellVoyager CQ1 (Yokogawa Electric). Additionally, we thank Yuki Mori and Kazuo Onishi from SCREEN Holdings for their valuable contributions to image acquisition and analysis using Cell3iMager duos2 (SCREEN Holdings).

## Supplementary material

Supplementary materials associated with this article can be found in the online version at doi:

