## Supplementary Material for "Development of a digital analysis system for a novel 3D culture-based colony formation to detect malignantly transformed cells in human cell-based therapeutic products"

$$y = \sum_{k=1}^{n\lambda} (n \cdot \frac{\lambda^k e^{-\lambda}}{k!}) \cdot (1 - (1 - x)^k)$$

| | |
|---|---|
| $x$ | Colony formation efficiency of reference cells (estimated in advance) |
| $y$ | Number of wells in which one or more colonies are detected |
| $\lambda$ | Percentage of cells distributed to each well |
| $n\lambda$ | Upper limit for the number of cells distributed in one well |
| $\frac{\lambda^k e^{-\lambda}}{k!}$ | Probability of containing k cells in one well (i.e. Poisson distribution) |
| $n \cdot \frac{\lambda^k e^{-\lambda}}{k!}$ | Number of wells containing k cells |
| $1 - x$ | Probability of one cell not forming a colony |
| $(1 - x)^k$ | Probability that all k cells seeded in one well do not form a colony |
| $1 - (1 - x)^k$ | Probability of formation of one or more colonies in one well |

**Supplementary Figure 1. The formula for calculating the colony-forming efficiency (CFE) of reference cells based on wells containing one or more detected colonies.**

The equation and accompanying table illustrate how the CFE is estimated from the number of wells that develop one or more colonies. Each parameter (x, y, λ, etc.) is defined, highlighting how cell distribution follows a Poisson process and how the probability of colony formation in each well is incorporated.

## a

**When seeding 840 wells with λ = 0.125, the following formula was applied based on the table below,**

$$y = \sum_{k=1}^{2} 840 \cdot \frac{\lambda^k e^{-\lambda}}{k!} \cdot (1 - (1 - x)^k)$$

**Before culture (when seeding)**

| Number of cells in one well | Probability distribution | Distribution of number of wells |
|---|---|---|
| 1 | 0.1103 | 93 |
| 2 | 0.0069 | 6 |
| 3 | 0.0003 | 0 |
| 4 | 0.00001 | 0 |
| 105 | 0.00000… | 0 |
| Total number of wells containing cells | | 99 |

**After culturing**

| Number of colonies in one well (*k*) | Distribution of the well number for each CFE (*x*) | | | | | | |
|---|---|---|---|---|---|---|---|
| | 100% | 90% | 80% | 70% | 60% | 50% | 40% |
| 1 | 93 | 83 | 74 | 65 | 56 | 46 | 37 |
| 2 | 6 | 6 | 6 | 5 | 5 | 4 | 4 |
| 3 | 0 | 0 | 0 | 0 | 0 | 0 | 0 |
| 4 | 0 | 0 | 0 | 0 | 0 | 0 | 0 |
| 105 | 0 | 0 | 0 | 0 | 0 | 0 | 0 |
| Total number of wells containing colonies (*y*) | 99 | 89 | 80 | 70 | 61 | 50 | 41 |

## b

**When seeding 420 wells with λ = 0.25, the following formula was applied based on the table below,**

$$y = \sum_{k=1}^{3} 420 \cdot \frac{\lambda^k e^{-\lambda}}{k!} \cdot (1 - (1 - x)^k)$$

**Before culture (when seeding)**

| Number of cells in one well | Probability distribution | Distribution of number of wells |
|---|---|---|
| 1 | 0.1947 | 82 |
| 2 | 0.0243 | 10 |
| 3 | 0.0020 | 1 |
| 4 | 0.0001 | 0 |
| 105 | 0.0000… | 0 |
| Total number of wells containing cells | | 93 |

**After culturing**

| Number of colonies in one well (*k*) | Distribution of the well number for each CFE (*x*) | | | | | | |
|---|---|---|---|---|---|---|---|
| | 100% | 90% | 80% | 70% | 60% | 50% | 40% |
| 1 | 82 | 74 | 65 | 57 | 49 | 41 | 33 |
| 2 | 10 | 10 | 10 | 9 | 9 | 8 | 7 |
| 3 | 1 | 1 | 1 | 1 | 1 | 1 | 1 |
| 4 | 0 | 0 | 0 | 0 | 0 | 0 | 0 |
| 105 | 0 | 0 | 0 | 0 | 0 | 0 | 0 |
| Total number of wells containing colonies (*y*) | 93 | 85 | 76 | 67 | 59 | 50 | 41 |

## c

**When seeding 240 wells with λ = 0.5, the following formula was applied based on the table below,**

$$y = \sum_{k=1}^{3} 240 \cdot \frac{\lambda^k e^{-\lambda}}{k!} \cdot (1 - (1 - x)^k)$$

**Before culture (when seeding)**

| Number of cells in one well | Probability distribution | Distribution of number of wells |
|---|---|---|
| 1 | 0.3033 | 73 |
| 2 | 0.0758 | 18 |
| 3 | 0.0126 | 3 |
| 4 | 0.0016 | 0 |
| 120 | 0.0000… | 0 |
| Total number of wells containing cells | | 94 |

**After culturing**

| Number of colonies in one well (*k*) | Distribution of the well number for each CFE (*x*) | | | | | | |
|---|---|---|---|---|---|---|---|
| | 100% | 90% | 80% | 70% | 60% | 50% | 40% |
| 1 | 73 | 66 | 58 | 51 | 44 | 36 | 29 |
| 2 | 18 | 18 | 17 | 17 | 15 | 14 | 12 |
| 3 | 3 | 3 | 3 | 3 | 3 | 3 | 2 |
| 4 | 0 | 0 | 0 | 0 | 0 | 0 | 0 |
| 105 | 0 | 0 | 0 | 0 | 0 | 0 | 0 |
| Total number of wells containing colonies (*y*) | 94 | 87 | 78 | 71 | 62 | 53 | 43 |

## d

**When seeding 120 wells with λ = 1.0, the following formula was applied based on the table below,**

$$y = \sum_{k=1}^{4} 120 \cdot \frac{\lambda^k e^{-\lambda}}{k!} \cdot (1 - (1 - x)^k)$$

**Before culture (when seeding)**

| Number of cells in one well | Probability distribution | Distribution of number of wells |
|---|---|---|
| 1 | 0.3679 | 44 |
| 2 | 0.1839 | 22 |
| 3 | 0.0613 | 7 |
| 4 | 0.0153 | 2 |
| 5 | 0.0031 | 0 |
| 120 | 0.0000… | 0 |
| Total number of wells containing cells | | 75 |

**After culturing**

| Number of colonies in one well (*k*) | Distribution of the well number for each CFE (*x*) | | | | | | |
|---|---|---|---|---|---|---|---|
| | 100% | 90% | 80% | 70% | 60% | 50% | 40% |
| 1 | 44 | 40 | 35 | 31 | 26 | 22 | 18 |
| 2 | 22 | 22 | 21 | 20 | 19 | 17 | 14 |
| 3 | 7 | 7 | 7 | 7 | 7 | 6 | 6 |
| 4 | 2 | 2 | 2 | 2 | 2 | 2 | 2 |
| 5 | 0 | 0 | 0 | 0 | 0 | 0 | 0 |
| 120 | 0 | 0 | 0 | 0 | 0 | 0 | 0 |
| Total number of wells containing colonies (*y*) | 75 | 71 | 65 | 60 | 54 | 47 | 40 |

**Supplementary Figure 2. Formulae for CFE calculation under multiple seeding conditions with different λ values.**

Shown here are the equations used to estimate the CFE of HeLa-green fluorescent protein (GFP) cells co-cultured with human mesenchymal stem cells (MSCs) at various average input levels (λ), including 0.125, 0.25, 0.5, and 1.0 cells/well, as presented in Figure 4b. The tables on the left illustrate the distribution of seeded cells per well according to the Poisson distribution, whereas the tables on the right show the theoretical distribution of colonies after a defined three-dimensional (3D) culture period, considering the probability of colony formation. For instance, when seeding cells at $\lambda = 0.25$ (cells/well) across 420 wells, it is reasonable to assume an upper limit of $k = 3$ (i.e., no more than three colonies will arise in a single well). Under this assumption, the number of wells containing at least one colony could be used to estimate the CFE of HeLa-GFP cells.

**a**

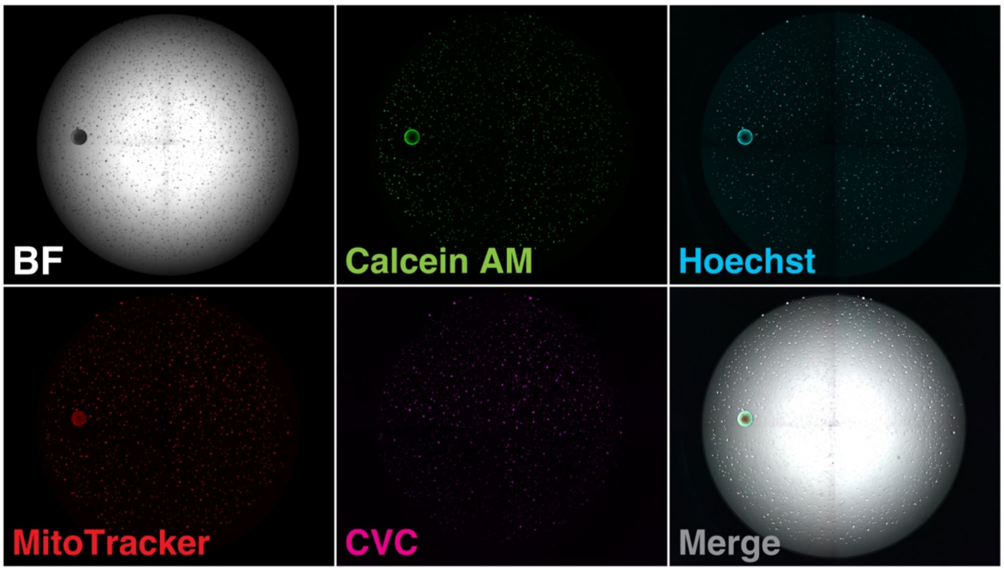


**b**

| Run | CFE (%) (~lower–upper) | Average CFE (%) (~lower–upper) |
|---|---|---|
| 1 | 35.1 (~30.5–41.1) | 33.0 (~27.7–40.0) |
| 2 | 33.1 (~28.3–39.2) | |
| 3 | 30.6 (~24.3–39.5) | |

**Supplementary Figure 3. Detection of HEK-293 cell-derived colonies after 3 weeks of co-culture using the D-LACF assay.**

(a) Representative images of a colony formed by a single HEK-293 cell after 3 weeks of co-culture with 17,000 normal human bone marrow-derived mesenchymal stromal/stem cells (MSCs) per well in 0.03% LA717-supplemented medium (200 µL/well). Before co-culture, both MSCs (PT-2501, Lot 0000471980; Lonza, Basel, Switzerland) and HEK-293 cells (ATCC CRL-1573, Lot 63777489; American Type Culture Collection, Manassas, VA, USA) were pre-labeled with CellVue Claret (CVC; Sigma-Aldrich, St.

Louis, MO, USA). Live-cell staining was performed at the end of the culture period by adding Hoechst 33342 (Dojindo Laboratories, Kumamoto, Japan), Calcein AM (Dojindo Laboratories), and MitoTracker Red CMXRos (Thermo Fisher Scientific, Waltham, MA, USA) to the culture medium. Images were acquired using a CellVoyager CQ1 (Yokogawa Electric, Tokyo, Japan). The inner diameter of each well was 6.4 mm. (b) Colony formation efficiency (CFE) of HEK-293 cells was assessed in three independent experimental runs. Each run involved seeding 300 wells with 0.2 HEK-293 cells and 17,000 MSCs per well, followed by a three-week culture period. CFEs were calculated based on the number of wells containing at least one colony. The lower and upper bounds of the 95% confidence interval (CI) for the CFE were determined based on the total number of HEK-293 cells spiked across wells in each run.

**Supplementary Movie. Real-time observation of a single HeLa-GFP cell forming a colony under co-culture with MSCs in medium supplemented with 0.03% LA717.**

This time-lapse movie (spanning 360 h at 1 h intervals) illustrates how the HeLa-GFP cell exhibits anchorage-independent growth, ultimately developing into a distinct colony, while the surrounding MSCs remain dispersed in a 3D environment maintained by LA717. The inner diameter of the well was 6.4 mm.

### Supplementary Table 1. Summary of CFE evaluation of HeLa-GFP cells at four different LA717 concentrations.

| LA717 | No. of spike cells (~lower–upper)[1] | Total number of wells seeded | Run | Total number of wells in which colonies were detected | $CFE_w$ (%)[2] (~lower–upper)[3] | Total number of detected colonies | $CFE_c$ (%)[4] (~lower–upper)[5] | Average $CFE_w$ (%)[6] (~lower–upper) | Average $CFE_c$ (%)[7] (~lower–upper) |
|---|---|---|---|---|---|---|---|---|---|
| 0.03% | 120 (~99–144) | 240 | 1 | 48 | 45.0 (~37.8–54.3) | 62 | 51.7 (~43.1–62.6) | 59.7 (~50.1–72.1) | 62.5 (~52.1–75.7) |
| | | | 2 | 68 | 67.1 (~56.3–81.0) | 82 | 68.3 (~56.9–82.8) | | |
| | | | 3 | 68 | 67.1 (~56.3–81.0) | 81 | 67.5 (~56.3–81.8) | | |
| 0.04% | | | 1 | 51 | 48.2 (~40.4–58.1) | 63 | 52.5 (~43.4–63.6) | 54.4 (~45.6–65.5) | 56.4 (~46.9–68.4) |
| | | | 2 | 60 | 58.0 (~48.6–69.9) | 75 | 62.5 (~52.1–75.8) | | |
| | | | 3 | 59 | 56.9 (~47.7–68.6) | 65 | 54.2 (~45.1–65.7) | | |
| 0.05% | | | 1 | 56 | 53.6 (~45.0–64.6) | 70 | 58.3 (~48.6–70.7) | 59.9 (~50.2–72.2) | 60.5 (~50.5–73.4) |
| | | | 2 | 76 | 76.7 (~64.3–92.5) | 91 | 75.8 (~63.2–91.9) | | |
| | | | 3 | 52 | 49.3 (~41.3–59.4) | 57 | 47.5 (~39.6–57.6) | | |
| 0.06% | | | 1 | 55 | 52.5 (~44.0–63.3) | 59 | 49.2 (~41.0–60.0) | 60.7 (~50.9–73.2) | 60.3 (~50.2–73.2) |
| | | | 2 | 64 | 62.5 (~52.4–75.4) | 78 | 65.0 (~54.2–78.8) | | |
| | | | 3 | 68 | 67.1 (~56.3–81.0) | 80 | 66.7 (~55.6–80.8) | | |

This table consolidates the colony formation data and the resulting colony-forming efficiency (CFE) calculations for HeLa-GFP cells co-cultured with mesenchymal stem/stromal cells (MSCs) at four LA717 concentrations (0.03–0.06%), as presented in Figure 4a.

[1] Lower and upper bounds of 95% confidence interval (CI) for total number of spiked cells.

[2] CFE calculated based on the number of wells containing at least one colony (newly proposed method in this study).

[3] The lower and upper bounds of the CFE were estimated using the formula shown in Supplementary Figure 1 and were calculated based on the 95% CI of the spiked cell number.

[4] CFE calculated based on the total number of colonies detected across all wells (conventional method).

[5] The lower and upper bounds of the CFE calculated using the conventional method were derived from the 95% CI of the spiked cell number.

[6] Average of three runs of $CFE_w$ and its lower and upper limits.

[7] Average of three runs of $CFE_c$ and its lower and upper limits.

### Supplementary Table 2. Summary of CFE evaluations for HeLa-GFP cells under four different seeding conditions (λ).

| λ[1] | No. of spike cells (~lower–upper)[2] | Total number of wells seeded | Run | Total number of wells in which colonies were detected | $CFE_w$ (%)[3] (~lower–upper)[4] | Total number of detected colonies | $CFE_c$ (%)[5] (~lower–upper)[6] | Average $CFE_w$ (%)[7] (~lower–upper) | Average $CFE_c$ (%)[8] (~lower–upper) |
|---|---|---|---|---|---|---|---|---|---|
| 0 | 0 | 60 | 1 | 0 | - | - | - | - | - |
| | | | 2 | 0 | - | - | - | | |
| | | | 3 | 0 | - | - | - | | |
| 0.125 | 105 (~85–128) | 840 | 1 | 76 | 76.1 (~55.9–94.1) | 78 | 74.3 (~60.9–91.8) | 78.6 (~57.7–97.1) | 76.2 (~62.5–94.1) |
| | | | 2 | 84 | 84.6 (~62.1–104.5) | 86 | 81.9 (~67.2–101.2) | | |
| | | | 3 | 75 | 75.1 (~55.1–92.8) | 76 | 72.4 (~59.4–89.4) | | |
| 0.25 | 105 (~85–128) | 420 | 1 | 52 | 49.3 (~41.3–59.4) | 53 | 50.5 (~41.4–62.4) | 70.6 (~53.3–86.8) | 71.8 (~58.9–88.6) |
| | | | 2 | 87 | 92.9 (~67.8–114.9) | 101 | 96.2 (~78.9–118.8) | | |
| | | | 3 | 67 | 69.6 (~50.8–86.1) | 72 | 68.6 (~56.3–84.7) | | |
| 0.5 | 120 (~99–144) | 240 | 1 | 71 | 70.6 (~59.2–85.2) | 82 | 68.3 (~56.9–82.8) | 78.7 (~66.0–95.0) | 75.8 (~63.2–91.9) |
| | | | 2 | 92 | 97.3 (~81.5–117.4) | 110 | 91.7 (~76.4–111.1) | | |
| | | | 3 | 69 | 68.3 (~57.3–82.4) | 81 | 67.5 (~56.3–81.8) | | |
| 1.0 | 120 (~99–144) | 120 | 1 | 57 | 65.1 (~54.9–78.5) | 78 | 65.0 (~54.2–78.8) | 75.5 (~63.6–91.0) | 72.0 (~60.0–87.2) |
| | | | 2 | 65 | 78.8 (~66.4–95.0) | 95 | 79.2 (~66.0–96.0) | | |
| | | | 3 | 67 | 82.6 (~69.6–99.5) | 86 | 71.7 (~59.7–86.9) | | |

This table consolidates the colony formation data and resulting CFE calculations for HeLa-GFP cells co-cultured with MSCs in 200 µL of the medium containing 0.03% LA717 at four different average input levels (λ), as presented in Figure 4b. Approximately 100 HeLa-GFP cells were evaluated for each condition.

[1] The expected average number of spiked cells per well at the time of seeding was derived from Poisson distribution.

[2] Lower and upper bounds of 95% CI for total number of spiked cells.

[3] CFE calculated based on the number of wells containing at least one colony (newly proposed method in this study).

[4] The lower and upper bounds of the CFE were estimated using the formula shown in Supplementary Figure 1 and were calculated based on the 95% CI of the spiked cell number.

[5] CFE calculated based on the total number of colonies detected across all wells (conventional method).

[6] Lower and upper bounds of the CFE derived from the conventional calculation method, based on 95% CI of the spiked cell number.

[7] Average of three runs of $CFE_w$ and its lower and upper limits.

[8] Average of three runs of $CFE_c$ and its lower and upper limits.

**Supplementary Table 3. Comparison between SACF and LACF assays.**

| Item | SACF | LACF |
|---|---|---|
| Preparation | +<br>Requires the preparation of multi-layer agar with varying concentrations of agarose | +++<br>A three-dimensional (3D) environment can be constructed using a single-layer setup when combined with low-adhesion vessels. |
| Temperature control | +<br>Requires strict temperature control to prevent gelation of the medium | +++<br>The medium stays liquid, which makes temperature control easy. |
| Imaging | +<br>Requires gel digestion and cell sedimentation before imaging | +++<br>Cells naturally settle at the bottom, enabling direct imaging. |
| Fixation | ++<br>Relatively easy; no strict temperature control needed | +<br>Requires strict temperature control to prevent polymer precipitation |
| Sample storage | +++<br>Storable at 4°C for over 1 month | ++<br>Storable at room temperature for up to 2 weeks |
| Culture condition flexibility | NA<br>Difficult to change agarose concentration or other conditions | ++<br>LA717 concentration can be modified based on the seeding density and cell type. |
| Time required for detecting a single HeLa cell-derived colony with MSC co-culture | ++<br>3–4 weeks | +++<br>2–3 weeks |
| Additional features | • Requires skilled manipulation<br>• Extensive information on usage history and applicable cell types is required. | • Culturing operations are easy.<br>• Cultured cells are easier to isolate and subculture. |

+++: Excellent; ++: Good; +: OK; NA: Not applicable.

LACF, liquid/low-molecular-weight agar colony formation; SACF, soft agar colony formation.

**Supplementary Table 4. Recommended requirements for imaging systems for colony detection.**

| Item | Recommended requirements | Notes |
|---|---|---|
| Plate format | 96-well plate | Standard reference format for whole-well analysis |
| Spatial resolution | ≤ 4 µm/pixel | Sufficient to resolve even small colonies of ~100 µm diameter |
| Camera resolution | ≥ 4 megapixels<br>(e.g., 2048 × 2048 pixels) | Allows some tiled images per well |
| Field of view (per shot) | ≥ 6.4 mm × 6.4 mm | Should cover at least one full well (tiling acceptable) |
| Sensor type | sCMOS recommended | High sensitivity, low noise, suitable for bright-field and fluorescence imaging |
| Multi-well imaging | Capable of continuous multi-well acquisition | Automated XY stage preferred |
| Tiling support | Required | Should include accurate image stitching and position correction |
| Focus control | Fixed focus or autofocus<br>(confocal preferred) | Basic focus functions are essential; confocal is a valuable addition. |
| Imaging modes | Bright-field and multi-channel fluorescence | Must support bright-field and multiple fluorescence channels |
| Image format | TIFF, PNG, JPG<br>(lossless compression preferred) | File formats suitable for quantitative analysis |
| Analysis application | • Basic analysis functions such as binarization-based segmentation and feature extraction of recognized objects<br>• Capable of continuous analysis across imaged wells | The application included in the imaging equipment that meet the above specifications is sufficient for analysis. |

sCMOS, scientific complementary metal-oxide semiconductor.

**Supplementary Table 5. Points to consider/troubleshooting for the D-LACF assay.**

| Item | Description |
|---|---|
| Preparation of quality control/positive control samples for the assay | • Use a quality control sample prepared by spiking a known quantity of reference (malignantly transformed) cells into the normal cell model.<br>• The normal cell model should preferably be the product cells themselves (same or different lot) or well-characterized diploid normal cells.<br>• Reference cells should be well-established and widely available from repositories (e.g., ATCC) and, although not required, are preferably relevant to the product type (e.g., glioma cell line for neural products). |
| Setting the target detection sensitivity and false-negative rate | • Determine a practical and clinically relevant detection sensitivity (e.g., 1 ppm) based on the number of transplanted cells and feasible assay scale (e.g., number of wells and plates).<br>• To calculate the required sample size (i.e., the number of wells to be seeded) for detecting transformed cells equivalent to the reference cells with a desired level of sensitivity, based on the colony formation rate of a single reference cell spiked into a fixed number of normal cells per well in a 96-well plate, it is necessary to predefine the acceptable false-negative rate. |
| Optimization of 3D culture conditions | • Adjust culture medium composition, seeding density, and LA717 concentration according to the characteristics of the product cells.<br>• If cell aggregation occurs at high densities, increasing LA717 concentration (up to ~0.06% w/v) may help suppress aggregation.<br>• Pre-labeling the samples with membrane dyes such as CellVue Claret or PKH26 can help distinguish aggregates from colonies. |
| Dealing with ambiguous colony identification | • When ambiguous cell clusters are frequently observed, isolate and recover the cells using a micropipette, and perform subculture to confirm proliferative capacity and malignancy potential.<br>• This step can also be useful for evaluating potential false positives. |
| Additional notes | • Prior to product testing, determine the required sample size (number of wells) based on performance testing using reference cells.<br>• Imaging threshold settings may vary depending on cell type and staining; a small-scale preliminary run is recommended to verify image analysis parameters. |
